# Topoisomerase I Essentiality, DnaA-independent Chromosomal Replication, and Transcription-Replication Conflict in *Escherichia coli*

**DOI:** 10.1101/2021.04.16.440247

**Authors:** J Krishna Leela, Nalini Raghunathan, J Gowrishankar

**Affiliations:** Laboratory of Bacterial Genetics, Centre for DNA Fingerprinting and Diagnostics, Hyderabad, India; Indian Institute of Science Education and Research Mohali, SAS Nagar, India

**Keywords:** DNA supercoiling, topoisomerase I, R-loops, constitutive stable DNA replication, transcription-replication conflict

## Abstract

Topoisomerase I (Topo I) of *Escherichia coli*, encoded by *topA*, acts to relax negative supercoils in DNA. Topo I deficiency results in hypernegative supercoiling, formation of transcription-associated RNA-DNA hybrids (R-loops), and DnaA- and *oriC*-independent constitutive stable DNA replication (cSDR), but some uncertainty persists as to whether *topA* is essential for viability in *E. coli* and related enterobacteria. Here we show that several *topA* alleles, including Δ*topA*, confer lethality in derivatives of wild-type *E. coli* strain MG1655. Viability in absence of Topo I was restored with two perturbations, neither of which reversed the hypernegative supercoiling phenotype: (i) in a reduced-genome strain MDS42, or (ii) by an RNA polymerase (RNAP) mutation *rpoB*35* that has been reported to alleviate the deleterious consequences of RNAP backtracking and transcription-replication conflicts. Four phenotypes related to cSDR were identified for *topA* mutants: (i) One of the *topA* alleles rescued Δ*dnaA* lethality; (ii) in *dnaA*^+^ derivatives, Topo I deficiency generated a characteristic copy number peak in the terminus region of the chromosome; (iii) *topA* was synthetically lethal with *rnhA* (encoding RNase HI, whose deficiency also confers cSDR); and (iv) *topA rnhA* synthetic lethality was itself rescued by Δ*dnaA*. We propose that the terminal lethal consequence of hypernegative DNA supercoiling in *E. coli topA* mutants is RNAP backtracking during transcription elongation and associated R-loop formation, which in turn lead to transcription-replication conflicts and to cSDR.

**Importance:** In all life forms, double helical DNA exists in a topologically supercoiled state. The enzymes DNA gyrase and topoisomerase I act, respectively, to introduce and to relax negative DNA supercoils in *Escherichia coli*. That gyrase deficiency leads to bacterial death is well established, but the essentiality of topoisomerase I for viability has been less certain. This study confirms that topoisomerase I is essential for *E. coli* viability, and suggests that in its absence aberrant chromosomal DNA replication and excessive transcription-replication conflicts occur that are responsible for lethality.

## Introduction

DNA in all cells is negatively supercoiled, and in bacteria such as *Escherichia coli* two enzymes gyrase and topoisomerase I (Topo I) ordinarily act oppositely to maintain the homeostasis of DNA superhelical density (reviewed in 1–4). DNA gyrase is a hetero-tetrameric enzyme (comprised of two subunits each of GyrA and GyrB proteins) that is ATP-dependent and introduces negative supercoils, whereas Topo I (encoded by *topA*) is an 865 amino acid-long monomer that catalyzes relaxation of supercoiled DNA in an energy-independent reaction. An important role for Topo I is in the dissipation of negative supercoils, in accord with the twin-domain supercoiling model (5, 6), that are generated behind RNA polymerase (RNAP) in the transcription elongation complex (TEC).

That gyrase deficiency leads to bacterial death is well established. On the other hand, the essentiality of Topo I for viability, in either *E. coli* or closely related bacteria such as *Salmonella enterica* and *Shigella flexneri*, is somewhat less certain (7–13). One difficulty has been that *topA* mutants rapidly accumulate suppressors which are often in the genes encoding the gyrase subunits (8–10, 13–17); and consistent with their opposing actions, gyrase and Topo I mutations can, in combination, partially cancel one another’s sickness or inviability (18, 19). Growth of *E. coli topA* mutants is also improved upon overexpression or amplification of genes encoding the topoisomerases III (13, 18, 20) or –IV (10, 16, 19).

Topo I deficiency is associated with an increased prevalence of R-loops (RNA-DNA hybrids) in the cells, which has been attributed to re-annealing of the 5’-end of nascent RNA into hyper-negatively supercoiled DNA behind the TEC under these conditions (21–23, reviewed in 4, 24, 25). Overexpression of RNase HI (encoded by *rnhA*), which degrades RNA in RNA-DNA hybrids, can alleviate some of the phenotypes of *topA* mutants (18, 22, 23, 26–28); and conversely, *topA rnhA* mutants exhibit exacerbated sickness (13, 26, 29). In principle, R-loops can exert toxicity by acting as road-blocks to subsequent transcription (30, 31) and to replication (32–34); and a third mechanism for toxicity is by serving as sites for initiation of aberrant chromosomal replication, as further outlined below. That R-loop formation is modulated by DNA supercoiling has been shown also in the CRISPR-Cas9 system (35), and in eukaryotic cells (36–38).

Recent evidence indicates that transcription-replication conflicts can themselves lead to increased formation of R-loops in the genome following RNAP backtracking at the sites of conflict (39–42, reviewed in 43–45). It has also been suggested that extended RNAP backtracking could be associated with R-loop formation from the 3’-end of the nascent RNA (40, 46).

R-loops are physiological initiators of ColE1 plasmid replication (47), but in addition their excessive occurrence (as in *rnhA* mutants) can lead to pathological initiation of chromosomal DNA replication in both bacteria (reviewed in 48–50) and eukaryotes (51). Such aberrant replication in bacteria is referred to as constitutive stable DNA replication (cSDR) since, unlike ordinary chromosomal replication which is initiated at *oriC* with the aid of the unstable protein DnaA, it continues long after inhibition of protein synthesis in the cells. cSDR can be identified biochemically as persistent DNA synthesis following addition of translational inhibitors such as chloramphenicol or spectinomycin.

cSDR can also be identified genetically as rescue of lethality associated with loss of DnaA function, which is a more stringent test of cSDR since it demonstrates the capability to duplicate the entire chromosome in the absence of *oriC*-initiated replication (48, 49). During its progression around the bacterial chromosome, such aberrant replication would be expected also to encounter (i) increased head-on conflicts with TECs on heavily transcribed genes (especially the *rrn* operons) that have evolved to be co-directional with *oriC*-initiated replisome progression, and (ii) increased arrest at *Ter* sites flanking the terminus region which are bound by the Tus protein (52, 53). The occurrence of cSDR in *rnhA* mutants has been established through both the biochemical and the genetic assays (48). The protein DksA, which participates in avoidance or resolution of transcription-replication conflicts (54, 55), is also required for viability of *rnhA dnaA* mutants (56). In recent work, Drolet’s group has shown by the biochemical assays that cSDR occurs in Topo I-deficient cells (28). Kornberg and coworkers had also shown earlier that specificity for replication initiation from *oriC in vitro* requires both RNase HI and Topo I (57, 58).

In this study, we have examined several *topA* insertion and deletion alleles for both their viability and their ability to rescue Δ*dnaA* lethality in *E. coli*. Our results indicate that *topA* null alleles are lethal in the wild-type strain MG1655 but that they are viable in MDS42, which is an engineered derivative lacking all prophages and transposable elements (59). The null mutants of MG1655 were viable with *rpoB*35*, which encodes an RNAP variant that has been reported to alleviate the deleterious effects of transcription-replication conflicts(40, 52, 60–65). Both in MDS42 and with *rpoB*35*, the viable Topo I-deficient derivatives continued to exhibit increased negative supercoiling. One *topA* allele could also rescue Δ*dnaA* lethality, providing genetic confirmation of cSDR in Topo I-deficient strains. We propose that bacterial lethality in absence of Topo I is caused by RNAP backtracking during transcription elongation and associated R-loop formation, which in turn then lead to transcription-replication conflicts and to cSDR.

## Results

### Description of topA insertion and deletion mutations and the assay to test for their viability

Three pairs of *topA* mutations were constructed on the *E. coli* chromosome by the recombineering approach of Datsenko and Wanner (66), each pair comprised of an FRT-flanked Kan^R^ element and the corresponding derivative with Flp recombinase-mediated site-specific excision of the Kan^R^ element to leave behind a “scar” of 27 in-frame codons; these are designated below by the suffixes “::Kan” and “::FRT”, respectively.

The three pairs of mutations represent the following (Fig. 1A): (i) deletion of all but the first codon and the last six codons (860-865) of the *topA* open-reading frame (Δ*topA*), that is, similar to the various gene knock-outs of the Keio collection (67); (ii) insertion beyond codon 480 in *topA* (*topA*-Ins480), this position being chosen because an earlier study had shown that a Tet^R^ insertion allele at this site was viable and associated with increased frequency of transposon precise excisions (11); and (iii) deletion beyond codon 480 until codon 860 in *topA* (*topA*-480Δ).

**Figure 1.**
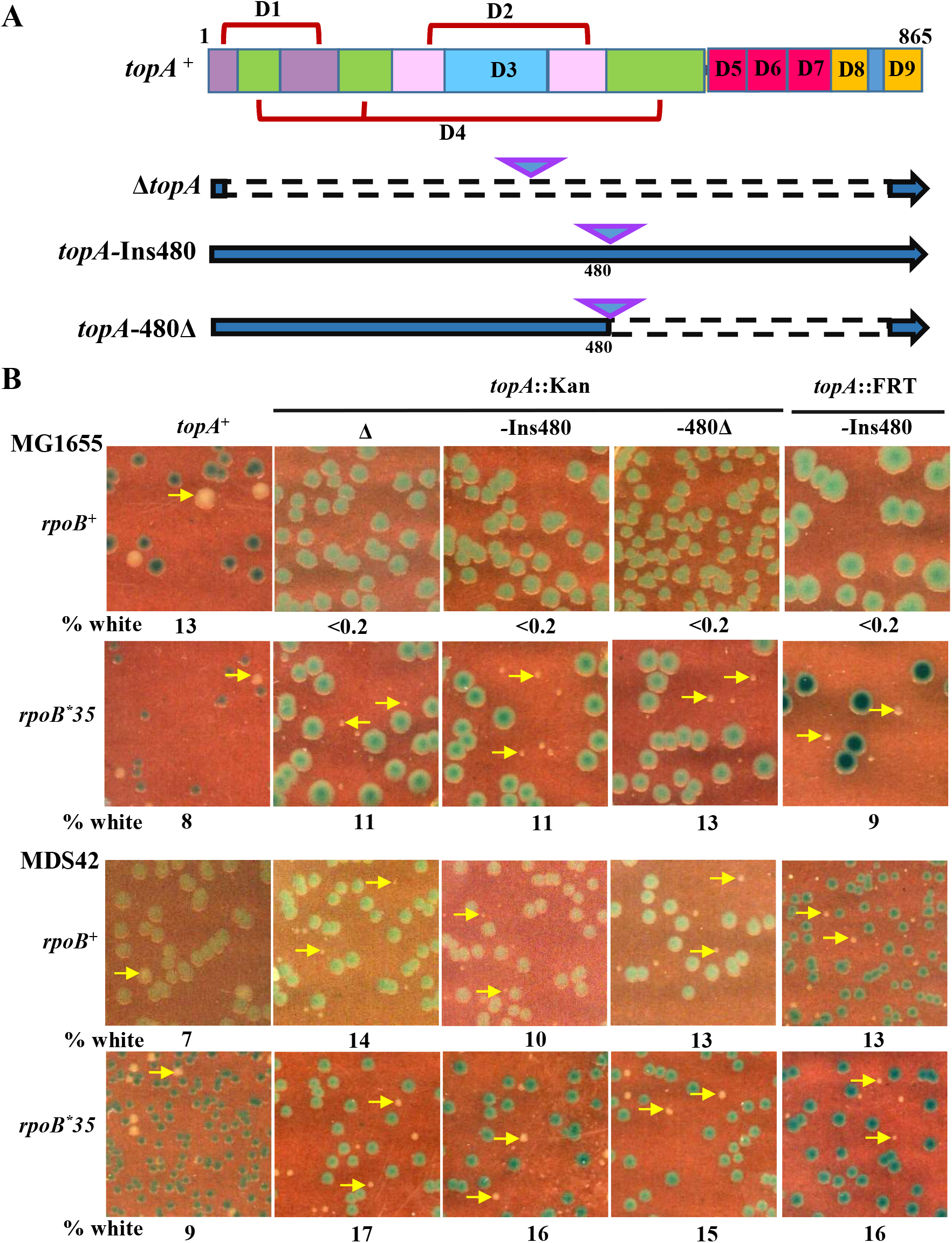
**(A)** Representations of *topA*^+^ ORF delineating the encoded protein’s domains D1 to D9 (adapted from 87, 88), and of the constructed *topA* alleles (three pairs) wherein the interrupted line segments represent deletions and each inverted triangle represents the pair comprising either Kan^R^ insertion (::Kan allele) or its FRT derivative (::FRT allele). **(B)** Blue-white screening assay on LB medium, of MG1655 or MDS42 strain derivatives with the *topA*^+^ shelter plasmid pHYD2390 and the different *topA* alleles as indicated on the top of each column; the *rpoB* allele status is indicated at left of each row. Examples of white colonies are marked by the yellow arrows. From left to right, strains used for the panels were pHYD2390 derivatives of: row 1, GJ13519, GJ15603, GJ15604, GJ15688 and GJ16921; row 2, GJ16703, GJ16813, GJ16814, GJ16815 and GJ16854; row 3, GJ12134, GJ16816, GJ16817, GJ16818 and GJ18977; and row 4, GJ16819, GJ16820, GJ16821, GJ16822 and GJ17777.

All the *topA* mutations were constructed and maintained in derivatives that were also Δ*lacZ* on the chromosome and carried a shelter plasmid derivative of a single-copy-number IncW replicon encoding trimethoprim (Tp)-resistance (68) with *topA*^+^ and *lacZ*^+^ genes. Since this plasmid’s segregation into daughter cells during cell division is not tightly controlled, around 10% of cells in a population are plasmid-free. Only provided that these latter cells are viable, they grow as white colonies on Tp-free medium supplemented with Xgal, whereas the plasmid-bearing cells grow as blue colonies on these plates. The appearance of white colonies which can be subsequently purified, therefore, is a demonstration of viability in absence of the *topA*^+^ shelter plasmid, and we have employed the similar blue-white screening assays earlier for tests of viability with other essential genes such as *rho, nusG, dnaA*, and *rne* (69–71).

### Viability of topA mutations in MDS42 and MG1655 rpoB*35

With the blue-white assay above, we found that none of the six *topA* alleles is viable in MG1655 (Fig. 1B). These observations are consistent with those of Stockum et al. (13) who also employed a similar approach to conclude that *topA* is essential in *E. coli*.

By the same blue-white assay, we could show that all the *topA* mutations are viable in strain MDS42 and in *rpoB*35* derivatives of both MG1655 and MDS42, on both LB and defined media (Fig. 1B and Supp. Fig. S1A); the growth of white colonies of the MDS42 *topA* derivatives was improved in presence of *rpoB*35*, on both media (Supp. Fig. S1B). MDS42 is a derivative of MG1655 with 14% of its genome (comprising all prophages and transposable elements) deleted (59), while *rpoB*35* is a mutation in RNAP that has been reported to render the enzyme less prone to backtracking or arrest and more accommodative of conflicts with replication (40, 52, 60–65).

In microscopy experiments (Supp. Fig. S2), cell size and morphology were unchanged with the *rpoB*35* mutation alone in both MG1655 and MDS42. Cells of the MDS42 *topA* mutant were filamented, and the filamentation was to a large extent suppressed in the *topA rpoB*35* derivative. The *topA rpoB*35* derivative of MG1655 was also moderately filamented.

Growth rate experiments in liquid cultures did not yield reliable data because of extended lag times and accumulation of suppressors in the *topA* derivatives. Suppressor accumulation has also been documented earlier for *topA* mutants by other workers (8, 13). Based on the observation that the “white” *topA* mutant clones in the blue-white screening assay grow to colonies of around 10^8^ cells in 48 hours, we have estimated a doubling time of around 100 min.

For reasons that are explained in the Discussion, we tested whether suppression of *topA* lethality by *rpoB*35* in MG1655 is abolished in absence of the UvrD DNA helicase in the cells. In the blue-white assay, viable colonies of MG1655 *rpoB*35 topA* were obtained even in a Δ*uvrD* background, indicating that the suppression is UvrD-independent (Supp. Fig. S3A).

### Rescue of topA lethality in MDS42 or by rpoB*35 is not because of reversal of hypernegative supercoiling

Lethality caused by loss of Topo I is associated with greatly elevated levels of negative supercoiling *in vivo*, and at least some suppressors of inviability, such as mutations in *gyrA* or *gyrB* and overexpression or amplification of genes encoding topoisomerases III or -IV, also confer reversal of the hypernegative supercoiling phenotype. To examine whether the viability of *topA* null mutants in MDS42 and with *rpoB*35* is correlated with reversal of hypernegative supercoiling, we determined their supercoiling status, by chloroquine-agarose gel electrophoresis (21, 72) of a reporter plasmid pACYC184 (73) in preparations made from the different strain derivatives.

The results, shown in Figure 2, indicate that (i) in MDS42, both *topA* mutations tested confer increased supercoiling (compare lanes 3-6 with lanes 1-2); (ii) *rpoB*35* does not alter supercoiling, in both the *topA*^+^ (compare within lane pairs 1 and 2, or 8 and 9) and *topA* derivatives (compare within lane pairs 3 and 4, or 5 and 6); and (iii) supercoiling levels are not different between the strain backgrounds of MG1655 and MDS42 for both *rpoB*^+^ (compare lanes 1 and 8) and the *rpoB*35* mutant (compare lanes 2 and 9). We conclude that when lethality conferred by loss of Topo I is suppressed by either genome-size reduction in MDS42 or *rpoB*35*, or by both perturbations together, there is no concomitant reduction in the hypernegative supercoiling status of DNA in these mutants.

**Figure 2.**
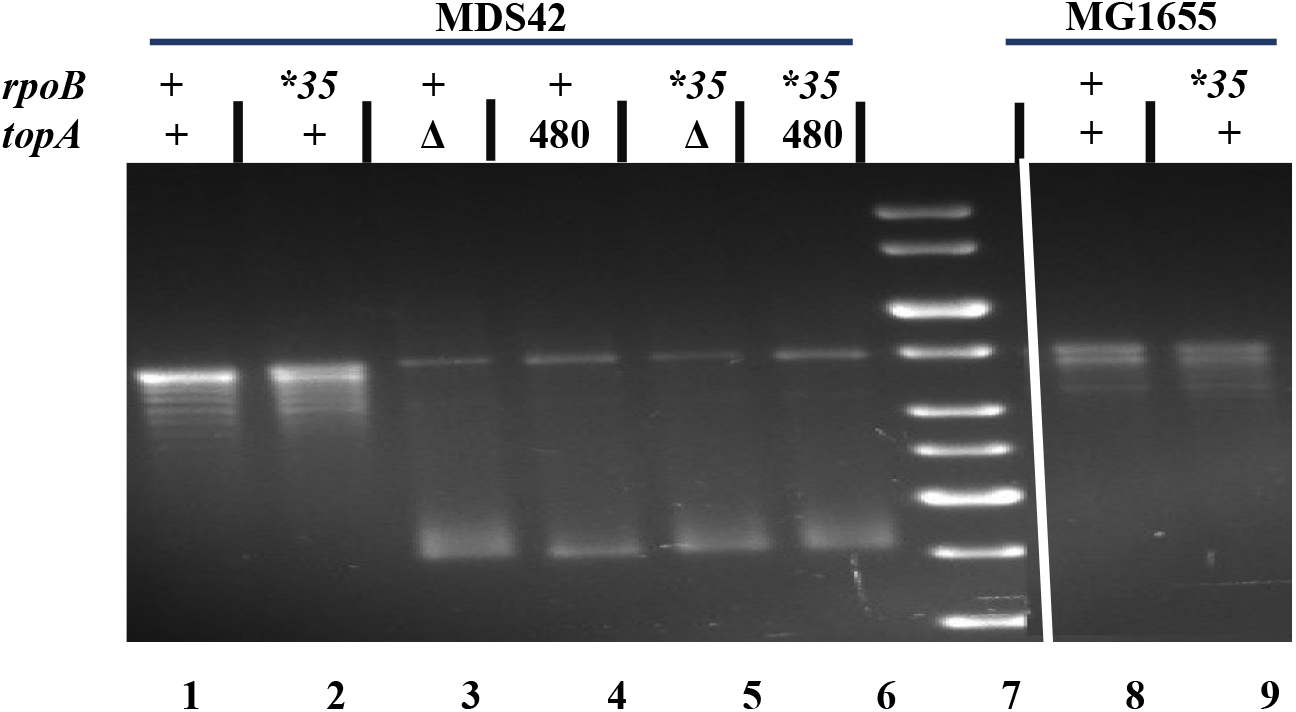
Supercoiling status of reporter plasmid pACYC184 in *topA*^+^ and *topA* derivatives, as determined by chloroquine-agarose gel electrophoresis. Genotypes at *topA* and *rpoB* loci are indicated on top of each lane; for *topA,* Δ and 480 refer, respectively, to Δ*topA*::FRT and *topA*-Ins480::FRT. Strains for different lanes were pACYC184 derivatives of: 1, GJ12134; 2, GJ16819; 3, 18976; 4, GJ18977; 5, GJ17776; 6, GJ17777; 8, GJ18601; and 9, GJ18910. Lane 7, DNA size standards.

### topA lethality in MG1655 is not rescued by ectopic expression of the R-loop helicase UvsW

Ectopic expression of the phage T4-encoded R-loop helicase UvsW (74) has previously been shown to rescue lethality associated with increased R-loop prevalence in several different *E. coli* mutants. The latter include strains with combined deficiency of RNases HI and HII (69), or of RNase HI and RecG (75), as also those with deletions of genes *rho* or *nusG* involved in factor-dependent transcription termination (69).

Since Topo I deficiency phenotypes are also associated with increased occurrence of intracellular R-loops and are partially suppressed by RNase HI overexpression (18, 21–23, 26–28), we employed the blue-white assays to examine whether UvsW expression (from a P_tac_-UvsW chromosomal construct, induced with IPTG) could rescue MG1655 *topA* lethality; an MG1655 Δ*rho* derivative (whose lethality is known to be rescued by UvsW) was chosen as control. The results indicate that UvsW expression does not confer viability to the MG1655 *topA* derivative, whereas it could do so to the Δ*rho* mutant (Supp. Fig. S3B). UvsW expression was associated with impaired growth of the *topA*^+^ blue colonies; this growth impairment was exemplified both by a marked decrease in plating efficiency and by occurrence of blue haloes around the colonies, suggestive of cell lysis. That UvsW expression is toxic to wild-type *E. coli* has been reported earlier (69).

### Rescue of ΔdnaA lethality by Topo I deficiency

As mentioned above, Topo I deficiency was earlier shown by a biochemical assay to confer cSDR, but whether it can rescue lethality associated with loss of DnaA function (that is, the genetic assay for cSDR) has not been determined. We adapted the blue-white assay to test whether any of the *topA* mutations can rescue lethality associated with loss of DnaA function, by constructing a Tp^R^ *lacZ*^+^ shelter plasmid that carried both *topA*^+^ and *dnaA*^+^. Three different *dnaA* alleles were used in these experiments: Δ*dnaA*::Kan (70), which is a Keio-style insertion-deletion that has removed all but the first codon and the last six codons of the 468-codon-long *dnaA* ORF; its FRT derivative, Δ*dnaA*::FRT (70); and *dnaA177* (76), whose DNA sequence determination revealed that it carries both a missense mutation in codon 267 (resulting in Thr-to-Ilv substitution) and an amber nonsense mutation in codon 450. The strains also carried Δ*tus* and *rpoB*35* mutations, which facilitate cSDR-directed chromosome duplication by overcoming the problems posed, respectively, by the *Ter* sites and by excessive head-on transcription-replication conflicts (52, 70, 77, 78).

Of the six *topA* mutations tested that had been shown above to be lethal in MG1655, one (*topA*-Ins480::FRT) was able to rescue lethality of Δ*dnaA*::FRT and of *dnaA177* at 30° on both minimal and LB media (Fig. 3A-B and Supp. Fig. S4A), while the others yielded no viable white colonies (Fig. 3A). Even with *topA*-Ins480::FRT, there was no rescue imposed by DnaA deficiency at 37° or 42° (Fig. 3B, see also row 5 in each of the panels of Supp. Fig. S5), nor were viable colonies recovered with the Δ*dnaA*::Kan allele (Fig. 3A). On the other hand, Δ*dnaA*::Kan lethality is rescued by other cSDR-provoking mutations such as *rnhA* or *dam* (data not shown).

**Figure 3.**
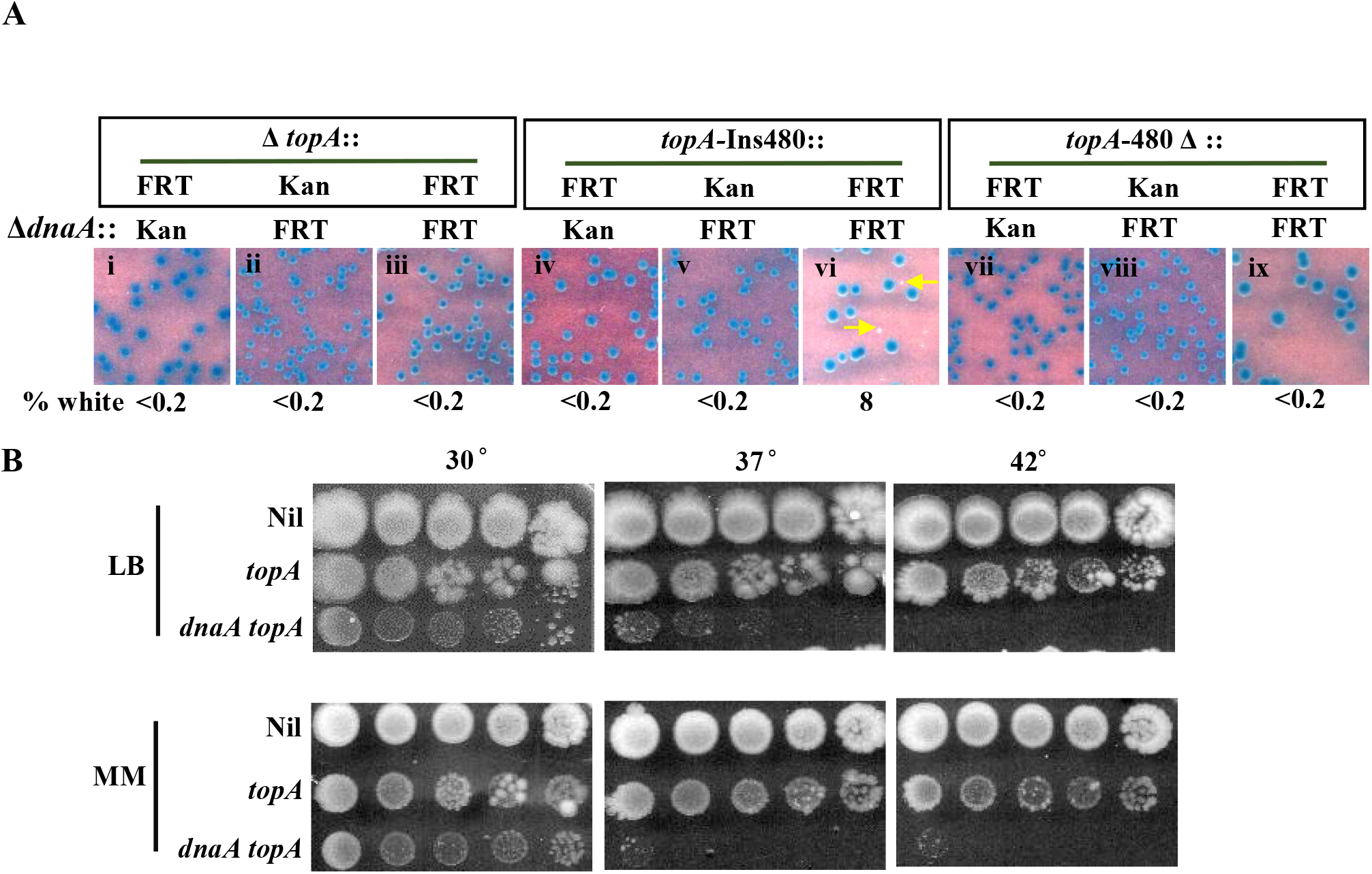
Suppression of Δ*dnaA* lethality by *topA*. *(A)* Blue-white screening assay at 30° on glucose-minimal A with *topA*^+^ *dnaA*^+^ shelter plasmid pHYD2390 in MG1655 Δ*dnaA* Δ*tus rpoB*35* derivatives carrying different *topA* alleles; the nature of Δ*dnaA* allele (::Kan or ::FRT) and of *topA* allele are shown on top of each panel. Examples of white colonies are marked by the yellow arrows. Strains employed for the different panels were pHYD2390 derivatives of: i, GJ17786; ii, GJ17790; iii, GJ18940; iv, GJ17787; v, GJ17791; vi, GJ18941; vii, GJ17788; viii, GJ17792; and ix, GJ18942. *(B)* Serial dilution-spotting on LB and glucose-minimal A (MM) at indicated temperatures of the following derivatives of MG1655 Δ*tus rpoB*35*: Nil, *topA*^+^ *dnaA*^+^ (GJ17784/pHYD2390); *topA*, *topA-*Ins480::FRT *dnaA*^+^ (GJ17784); and *topA dnaA*, *topA-*Ins480::FRT Δ*dnaA*::FRT, that is, white colony from panel vi of Figure 4A (GJ18941).

Two distinct and interesting interpretations are suggested from these data: (i) that unlike the other five *topA* alleles, *topA*-Ins480::FRT might possess a low level of DNA relaxation activity (since it encodes a full-length polypeptide with just a 27-amino acid linker inserted between residues 480 and 481 of Topo I) which is not sufficient for viability *per se* in MG1655 but nonetheless is necessary for viability in the derivatives whose sole source of chromosome duplication is cSDR; and (ii) that expression of the essential *dnaN* gene immediately downstream of, and in the same operon as, *dnaA* is achieved from a fortuitous outward reading promoter in the Kan^R^ element of the Δ*dnaA*::Kan allele, but that this promoter is rendered inactive under TopoI-deficient conditions.

Notwithstanding these unusual features, our data clearly establish that cSDR in a Topo I-deficient derivative can act to rescue the lethality associated with total absence of DnaA in the cells. This viability is contingent on absence of the Tus protein (Supp. Fig. S4B). On the other hand, the DinG helicase, which has been shown to be needed for Δ*dnaA* viability of RecG- or Dam-deficient cells (70), was not so required in the Topo I-deficient strain, nor did its absence impede viability of the *topA* mutant in the *dnaA*^+^ derivative (Supp. Fig. S4C).

### Copy number analysis of different chromosomal regions in topA mutants

In an exponentially growing population of bacterial cells, DnaA-initiated replication imposes a bidirectional gradient of copy number for different regions of the circular chromosome, with the peak near *oriC* and trough in the terminus region (53). If a *dnaA*^+^ strain also suffers a perturbation that activates cSDR (such as deficiency of RNase HI, RecG, Dam, or multiple DNA exonucleases), the DNA copy number pattern is characterized by superposition of a “mid-terminus peak” on the bidirectional gradient described above (52, 70, 77–82). We have earlier proposed that the mid-terminus peak represents a population aggregate of replication forks progressing from stochastically firing cSDR origins that are widely distributed across the genome (53, 70), although other groups have suggested that it represents a discrete origin of replication (52, 77–79), or occurrence of over-replication when oppositely directed forks converge at the terminus (80, 81).

Drolet and coworkers have shown earlier that Topo I-deficient mutants exhibit the mid-terminus peak (19), but their strains also carried additional genetic changes such as a *gyrB* (Ts) mutation and amplification of the genes encoding subunits of topoisomerase IV. For our DNA copy number analysis studies, we used *dnaA*^+^ strains of the MDS42 background without or with *rpoB*35* and additionally with the *topA*-Ins480::FRT mutation (that is, the allele associated with rescue of Δ*dnaA* inviability).

The whole genome sequence reads obtained from each of the strains were aligned to the MDS42 reference sequence, and normalized read counts for the different chromosomal regions were determined. No suppressor mutation in any of the candidate genes was identified in the *topA* mutants, while presence of the *topA* mutation itself and of the CAC-to-CAA codon change (His-to-Gln substitution) associated with the *rpoB*35* allele (60) was confirmed in each of the relevant strains.

The parental (*topA*^+^ *rpoB*^+^) strain exhibited the expected bidirectional copy number gradient from *oriC* to *Ter* (Fig. 4, panel i), which was also largely preserved in its *rpoB*35* derivative (Fig. 4, panel ii). The *topA* mutant derivatives of both these strains showed distinct mid-terminus peaks superimposed on the *oriC*-to-*Ter* gradient (Fig. 4, panels iii and iv, respectively).

**Figure 4.**
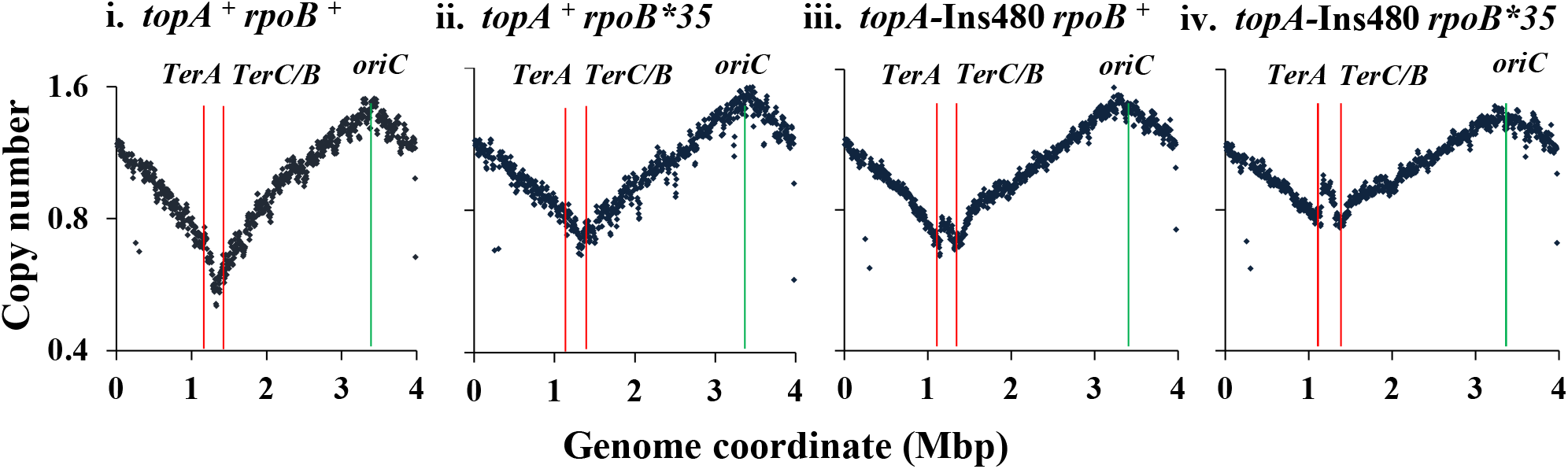
Copy number analysis by deep sequencing in *topA* mutant derivatives of MDS42. In panels i to iv, relative copy numbers have been plotted as semi-log graphs for overlapping 10-kb intervals across the genome (relevant genotype of strain indicated on top of each panel); positions of *oriC, TerA* and *TerC/B* are marked, and the gap at around 0.3 Mbp in each of the plots corresponds to the *argF-lac* deletion present in the strains. Strains displayed in the different panels are: i, GJ12134; ii, GJ16819; iii, GJ18977; and iv, GJ17777.

These results therefore confirm that a *topA* mutation capable of conferring Δ*dnaA* viability is associated with a mid-terminus peak of DNA copy number in *dnaA*^+^ derivatives.

### Mutual suppression between lethal mutations: loss of DnaA suppresses topA-rnhA synthetic lethality

Deficiency of either Topo I or RNase HI is associated with cSDR, and Stockum et al. (13) as well as Drolet and coworkers (26, 29) have earlier reported lethality or aggravated sickness in the double-deficient strains. We too found in this study that introduction of the *rnhA* mutation into otherwise viable *topA* derivatives (that is, in the MG1655-derived strain with *rpoB*35* and Δ*tus* mutations) confers synthetic lethality; the two *topA* alleles tested were the FRT derivatives of *topA*-Ins480 and Δ*topA* (Fig. 5, compare panels i-ii for former, and iv-v for latter). The lethalities were rescued in presence of Δ*dnaA* (Fig. 5, panels iii and vi, respectively), indicating that two otherwise lethal mutant combinations (*topA rnhA* and *dnaA*) could mutually suppress one another. Robust viability of the triple mutants was observed on both rich and defined media at 30° and 37°, less so at 42° (Supp. Fig. S5).

**Figure 5.**
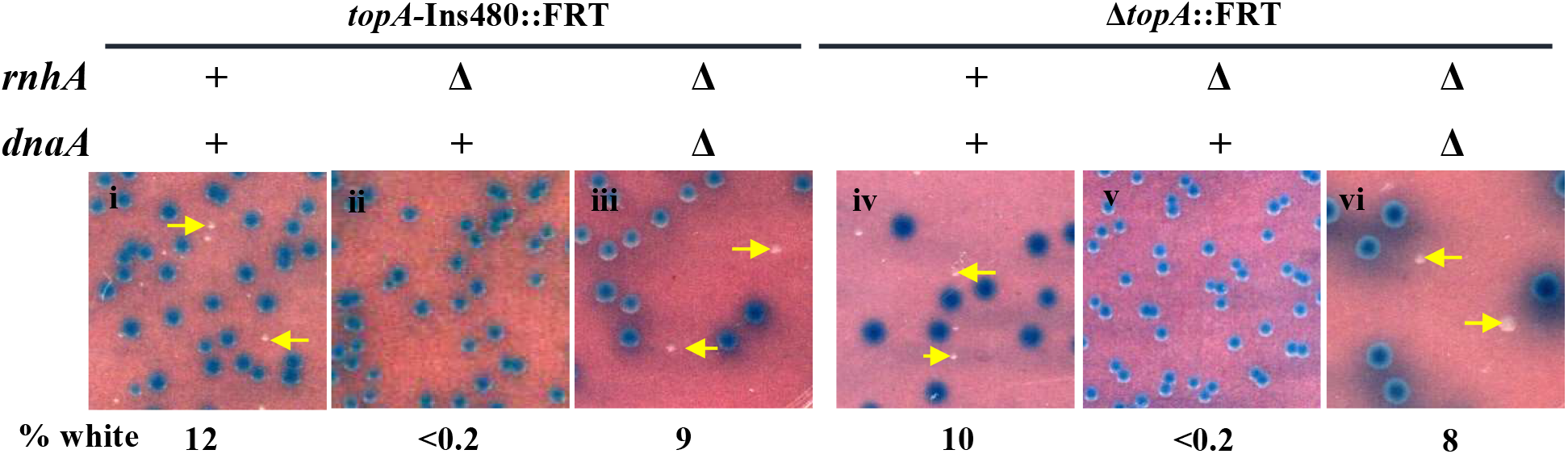
Synthetic *topA rnhA* lethality, suppressed by Δ*dnaA*. Blue-white screening assay at 30° on glucose-minimal A with *topA*^+^ *dnaA*^+^ shelter plasmid pHYD2390 in MG1655 Δ*tus rpoB*35* derivatives carrying different alleles of *topA*, *rnhA* and *dnaA* as indicated on top of each panel. Strains employed for the different panels were pHYD2390 derivatives of: i, GJ17784; ii, GJ19609; iii, GJ18951; iv, GJ17783; v, GJ19608; and vi, GJ18983.

We performed PCR experiments to confirm that the chromosomal *topA* locus in each of the viable triple mutant *topA rnhA dnaA* derivatives was indeed disrupted (and had not, for example, become *topA*^+^ by gene conversion from the wild-type allele on the shelter plasmid). Two primer pairs were used simultaneously to distinguish between the *topA*^+^, Δ*topA*::FRT, and *topA*-Ins480::FRT alleles, that yielded amplicons of 500, 328, and 581 bp, respectively (Supp. Fig. S6A-B). The results established that the signatures for the *topA*-Ins480 and Δ*topA* mutations were present, and that for *topA*^+^ was absent, in the triple mutant strains (Supp. Fig. S6B, lanes 5 and 7, respectively).

As discussed below, these results suggest that it is excessive chromosomal replication which kills cells doubly defective for Topo I and RNase HI.

## Discussion

The enzymes Topo I and DNA gyrase act to maintain the homeostatic balance of DNA negative supercoiling in *E. coli*. Topo I relaxes negative supercoils, especially those occurring behind RNAP during transcription elongation, and thus prevents the nascent transcript from re-annealing with the template DNA strand to form R-loops.

In this study, we have confirmed that Topo I-deficient *E. coli* mutants are inviable, and furthermore have identified two novel means by which the lethality can be independently and additively suppressed: (i) by deletion of the non-essential 14% of the genome comprising prophages and transposable elements, as in the strain MDS42; and (ii) by the *rpoB*35* mutation encoding an altered RNAP which has earlier been variously described (not mutually exclusive) to mimic the transcriptional effects of ppGpp (60), to reduce RNAP backtracking (40), and to mitigate the effects of transcription-replication conflicts by destabilizing the TEC (40, 52, 62, 63, 65). Neither of the suppressors acts by reversing hypernegative supercoiling in the *topA* mutants. We have also shown that Topo I deficiency, in the presence of additional *rpoB*35* and Δ*tus* mutations, can rescue Δ*dnaA* lethality, thereby providing genetic confirmation for occurrence of cSDR in the Topo I-deficient derivatives.

### rpoB*35 and RNAP backtracking

As mentioned above, several workers have provided evidence that the *rpoB*35*-encoded substitution in RNAP destabilizes the TEC in vitro (40, 62) and protects against transcription-replication conflicts in vivo (65), including during cSDR (52, 70, 77, 78). Trautinger and Lloyd (61) have reported that *rpoB*35* suppresses the Ts phenotype of *greA greB* double mutants and the UV-sensitivity of an *mfd* mutant, which they interpret as evidence that it may function by preventing backtracking, thus facilitating dissociation of stalled TECs. Likewise, *rpoB*35* also suppresses *rep uvrD* lethality, which has been ascribed to direct reduction of replicative barriers posed by TECs under these conditions (63).

On the other hand, there is one report from the Nudler group that RpoB*35-substituted RNAP exhibits increased backtracking in vitro in presence of UvrD (64). This property of RpoB*35-RNAP appears to be strictly UvrD-dependent, and the same group has shown in other studies (40) that relative to wild-type RNAP, the RpoB*35 enzyme is resistant to pausing and backtracking.

It is therefore reasonable to conclude that the RpoB*35 enzyme is in general more resistant than wild-type RNAP to arrest and backtracking during transcription elongation, except perhaps in the specific context when a high concentration of UvrD dimers occurs following DNA damage. Our finding in this study, that the suppression of *topA* lethality by *rpoB*35* is UvrD-independent (Supp. Fig. S3A), is noteworthy in this context.

### Mechanism of lethality in Topo I-deficient strains

The fact that *rpoB*35* restores growth to MG1655 in absence of Topo I without affecting the hypernegative supercoiling status of the mutants suggests that it is the downstream consequences of increased negative supercoiling, namely RNAP backtracking and impairment of TEC progression leading to transcription-replication conflicts, which are responsible for *topA* lethality. Pathological R-loop formation is also expected to be an important feature at the arrested TECs, but whether it precedes or follows RNAP backtracking remains to be determined. In the *topA* mutant, *rpoB*35* would also relieve the sickness during cSDR engendered by transcription-replication conflicts especially at the *rrn* operons.

To explain *topA* viability in MDS42, we propose that the regions of the genome that are deleted in this strain (comprising prophages and transposable elements) are preferentially enriched for sites of R-loop formation, TEC arrest and transcription-replication conflict. Loss of the proteins Rho or NusG, that are involved in factor-dependent transcription termination and reportedly in R-loop avoidance (31, 69, 83–85), is also better tolerated in MDS42 than in MG1655, and especially so in presence of *rpoB*35* (65, 69, 86).

Finally, why does ectopic expression of the R-loop helicase UvsW not rescue *topA* lethality, although it can rescue the lethalities associated with loss of RNase H enzymes, RecG, Rho or NusG (69, 75)? One possibility is that R-loop formation in Topo I-deficient strains is a consequence, and not cause, of RNAP backtracking and arrest, so that R-loop removal per se would not mitigate the primary problem. An alternative possibility is that Topo I itself is required to relax the negative supercoils arising from UvsW’s helicase action on R-loops, and hence that UvsW is unable to act efficiently to unwind R-loops specifically in the *topA* mutants. The fact that RNase HI overexpression can suppress *topA* sickness phenotypes (26, 27) lends support to the second model.

### Topo I deficiency and cSDR

Drolet and coworkers had provided biochemical evidence for cSDR in Topo I-deficient cultures (28), which is presumably initiated from R-loops in these cells; our data establish that such cSDR is sufficient to sustain viability in absence of DnaA, in derivatives carrying Δ*tus* and *rpoB*35* mutations. The latter two mutations are expected to facilitate the completion of replication of the circular chromosome by forks initiated from site(s) other than *oriC* (52, 70, 77, 78). Our data also support the earlier suggestion (70) that incubation at 30° is more permissive than that at 37° or 42° for rescue by cSDR of Δ*dnaA* lethality.

Of the six different *topA* mutations that were inviable in MG1655-derived strains, it was only the *topA*-Ins480::FRT allele that could confer viability to the Δ*dnaA* derivatives. As explained above, this mutation generates a modified version of Topo I in which a 27 amino acid-linker is inserted between residues 480 and 481 of the polypeptide. From the Topo I monomer crystal structure (87, 88), it is expected that the linker is situated at or near the junction between residues that comprise domain D2 and those that comprise domain D4; it is therefore possible that the linker may allow for (the albeit inefficient) folding of the polypeptide to yield a correct tertiary structure. The residual Topo I activity of this protein might be needed for proper chromosome segregation after cSDR in the Δ*dnaA* mutants (20).

### oriC-initiated replication contributes to topA-rnhA synthetic lethality

We have shown that although the Δ*topA* and Δ*rnhA* combination is synthetically lethal, the Δ*topA* Δ*rnhA* Δ*dnaA* mutant is viable. Thus, *oriC*- initiated replication is a contributor to *topA rnhA* toxicity, which suggests that it is excessive replication (sum of that from *oriC* and cSDR, the latter contributed additively by both *rnhA* and *topA*) which confers toxicity. These results are in agreement with those from Drolet and coworkers (20, 29), who had earlier reported that mutations affecting either replication from *oriC* or replication restart functions can alleviate the sickness of cells deficient for both Topo I and RNase HI activities.

## Materials and Methods

### Growth media, bacterial strains and plasmids

The rich and defined growth media were, respectively, LB and minimal A with 0.2% glucose (89) and, unless otherwise indicated, the growth temperature was 37°. Supplementation with Xgal and with antibiotics ampicillin, kanamycin (Kan), tetracycline (Tet), chloramphenicol (Cm), and trimethoprim (Tp) were at the concentrations described earlier (90). Isopropyl-β-D-thiogalactoside (IPTG) was added at the indicated concentrations. *E. coli* strains used are listed in Table S1, in the supplemental material.

Plasmids described earlier include pMU575 (Tp^R^, single-copy-number vector with *lacZ*^+^) (68); pACYC184 (Tet^R^ Cm^R^, p15A replicon) (73); pHYD2388 (70) and pHYD2411 (69) (Tp^R^, pMU575 derivatives with, respectively, *dnaA*^+^ and *rho*^+^); and pKD13, pKD46 and pCP20 described by Datsenko and Wanner (66) for recombineering experiments and Flp-recombinase catalyzed excision between a pair of FRT sites. The construction of two derivatives of plasmid pMU575 is described in the supplemental material: pHYD2382, carrying *topA*^+^, and pHYD2390, carrying *topA*^+^ *dnaA*^+^.

### Blue-white screening assays

To determine lethality or viability of strains with chromosomal *topA* or *dnaA* mutations, derivatives carrying the shelter plasmids pHYD2382 (*topA*^+^) or pHYD2390 (*topA*^+^ *dnaA*^+^) were grown overnight in Tp-supplemented medium, subcultured into medium without Tp for growth to mid-exponential phase, and plated at suitable dilutions on Xgal plates without Tp. The percentage of white colonies to total was determined (minimum of 500 colonies counted), and representative images were captured. *Plasmid supercoiling assays.* Strains carrying plasmid pACYC184 were grown in LB to mid-exponential phase, and plasmid preparations were made with the aid of a commercial kit. Plasmid supercoiling status in each of the preparations was determined essentially as described (72), following electrophoresis on 1% agarose gel with 5 µg/ml chloroquine at 3 V/cm for 17 hr.

### Copy number analysis by deep sequencing

Copy number determinations of the various genomic regions were performed essentially as described (70). Genomic DNA was extracted by the phenol-chloroform method from cultures grown in LB to mid-exponential phase, and single-end deep sequencing was performed on Illumina platforms to achieve around 100-fold coverage for each preparation. Sequence reads were aligned to the MDS42 reference genome (Accession number NC_020518.1), and copy numbers were then determined by a moving-average method after normalization of the base read count for each region to the aggregate of aligned base read counts for that culture.

### Other methods

Procedures were as described for P1 transduction (91) and for recombinant DNA manipulations, PCR, and transformation (92). Different chromosomal *topA* mutations were generated by recombineering (66) as described in the supplemental material. For microscopy experiments, cells from cultures grown in LB to mid-exponential phase were immobilized on 1% agarose pads and visualized by differential interference contrast imaging with the aid of a Zeiss Axio Imager Z2 microscope.

### Data availability

Genome sequence data from this work have been submitted under Accession number PRJNA670792 and are available for public access at https://www.ncbi.nlm.nih.gov/Traces/study/?acc=PRJNA670792.

## Supplemental material

Supplemental material is provided as a PDF file “Supplemental File 1”.

## Acknowledgements

We thank Jillella Mallikarjun for sequencing the *dnaA177* allele; Aswin Seshasayee and T V Reshma for help with deep sequencing; V Balaji for assistance with microscopy; and the COE team members for advice and discussions.

This work was supported by Government of India funds from (i) DBT Centre of Excellence (COE) project for Microbial Biology – Phase 2, (ii) SERB project CRG/2018/000348, and (iii) DBT project BT/PR34340/BRB/10/1815/2019. JG was recipient of the J C Bose fellowship and INSA Senior Scientist award.

We declare that there are no conflicts of interest.

## Supplemental Material for

### Supplementary Methods

#### Construction of shelter plasmids pHYD2382 and pHYD2390

The pair of primers 5’-CCAAA*CTGCAG*TCGTGCTATAGCGCCTGT-3’ and 5’-TTTGTT*AAGCTT*AACCTGACAGAATTAAAGG -3’ (*PstI* and *HindIII* sites in them, respectively, italicized) were used with MG1655 genomic DNA to amplify the *topA*^+^ gene by PCR, and the product was digested with *PstI* and *HindIII* before being cloned into the corresponding sites of plasmid vector pMU575, to generate the *topA*^+^ shelter plasmid pHYD2382. The two primers are designated, respectively, as b1 and b2 in Supplementary Figure S7.

For constructing plasmid pHYD2390 (*topA*^+^ *dnaA*^+^ shelter plasmid), the DNA fragment present in plasmid pHYD2388 (1) carrying the *Salmonella enterica dnaA*^+^ gene was sub-cloned *via* an intermediate plasmid vector into the *XbaI-KpnI* sites of plasmid pHYD2382.

#### Generation of topA mutations by recombineering

The Datsenko and Wanner method (2), with plasmids pKD13 and pKD46, was used for generating chromosomal *topA* mutations by recombineering in derivatives carrying the shelter plasmids pHYD2382 or pHYD2390. The primer pairs used for PCR with pKD13 template were: for Δ*topA*::Kan, 5’- TCAACGTGCGACGCATTCCTGGAAGAATCAAATTAGGTAAGGTGAATATGATTCCGGGGATCCGTCGACC-3’ and 5’- TGTTTATAAAAACCTGACAGAATTAAAGGTTATTTTTTTCCTTCAACCCATGTAGGCTGGAGCTGCTTCG-3’ (designated primer Y); for *topA-*Ins480::Kan, 5’- CCAGCACTTTACCAAGCCGCCAGCCCGTTTCAGTGAAGCAATTCCGGGGATCCGTCGACC-3’ (designated primer X) and 5’- GACGACCGATACCGCGTTTTTCCAGCTCTTTAACCAGCGATGTAGGCTGGAGCTGCTTCG-3’; and for *topA-*480Δ, primers X and Y.

Given that the recombineering in these derivatives could have occurred on either the chromosome or the shelter plasmid, we employed simple genetic tests on several independent clones to classify them into the two categories. The three *topA*::FRT alleles on the chromosome were generated with the aid of plasmid pCP20, as described (2).

---

**Table S1.**
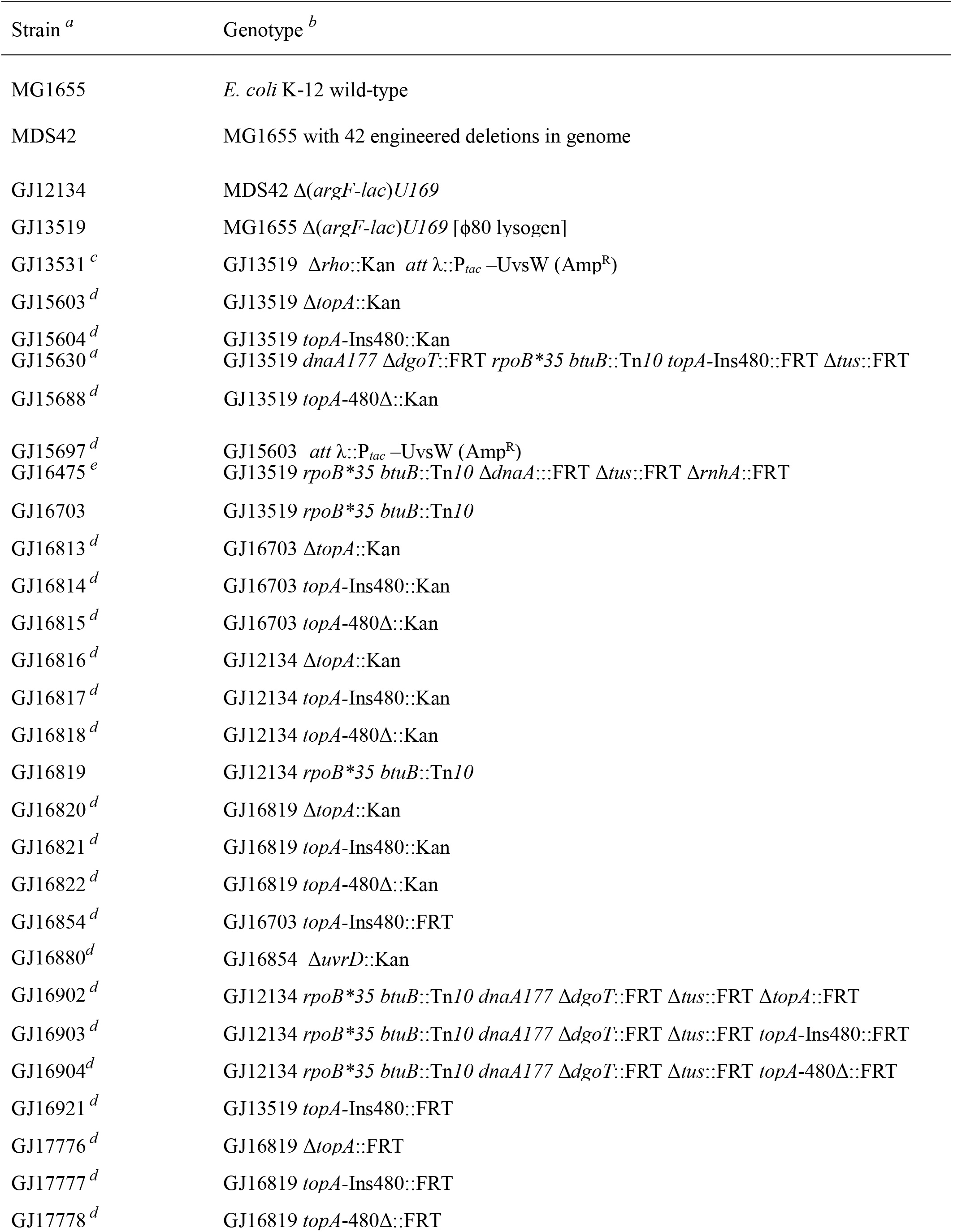
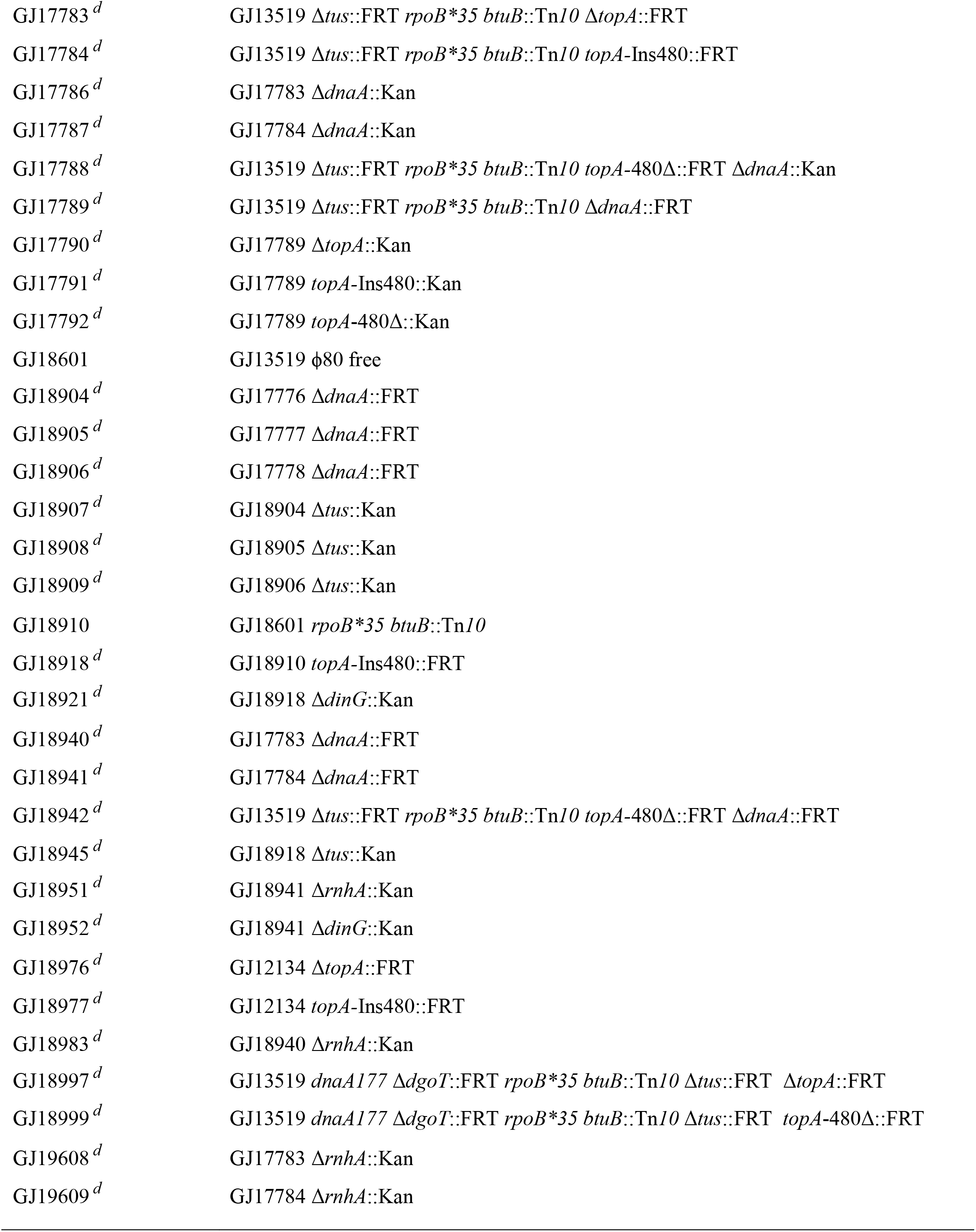
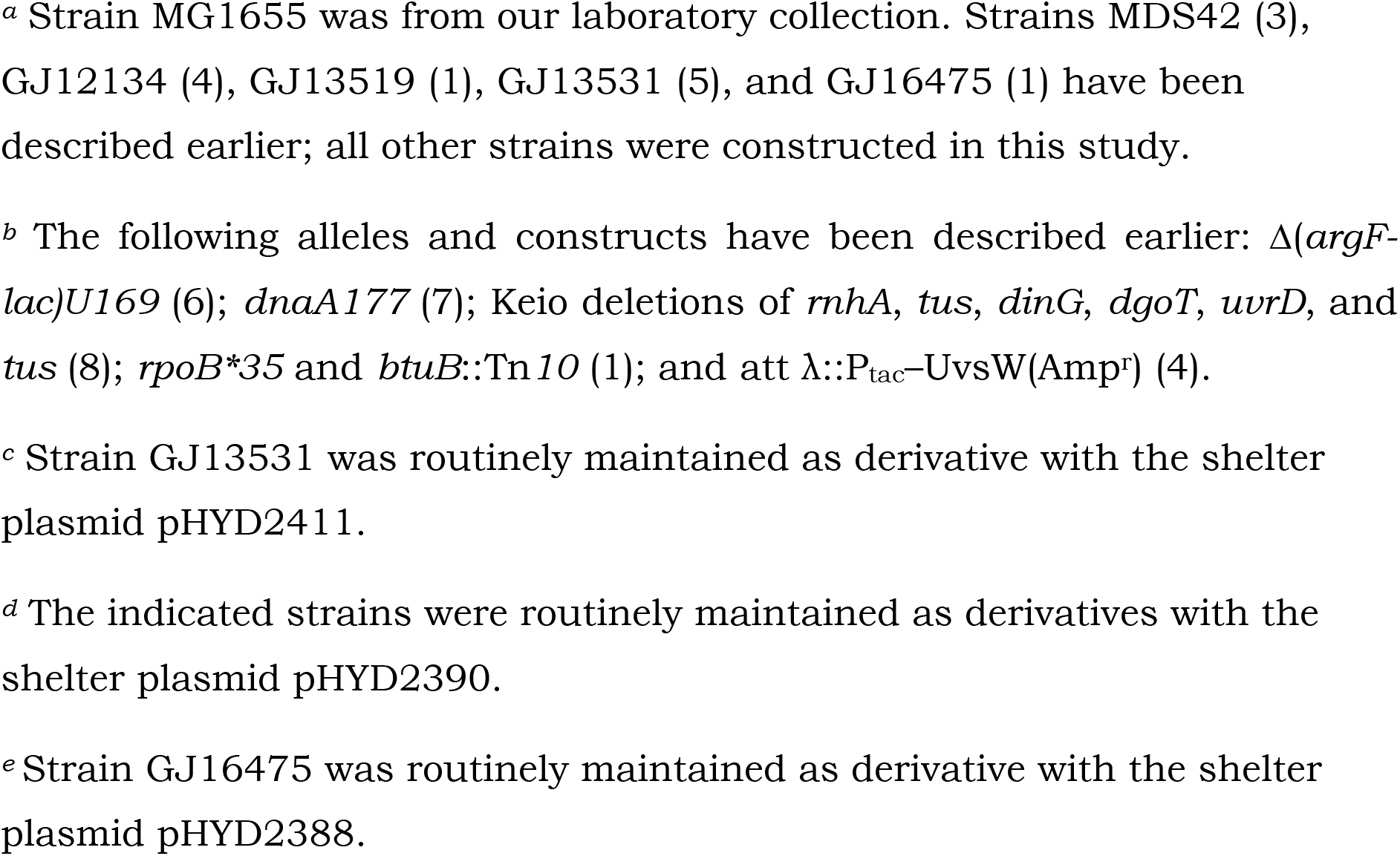
List of E. coli strains

**Supplementary Figure S1.**
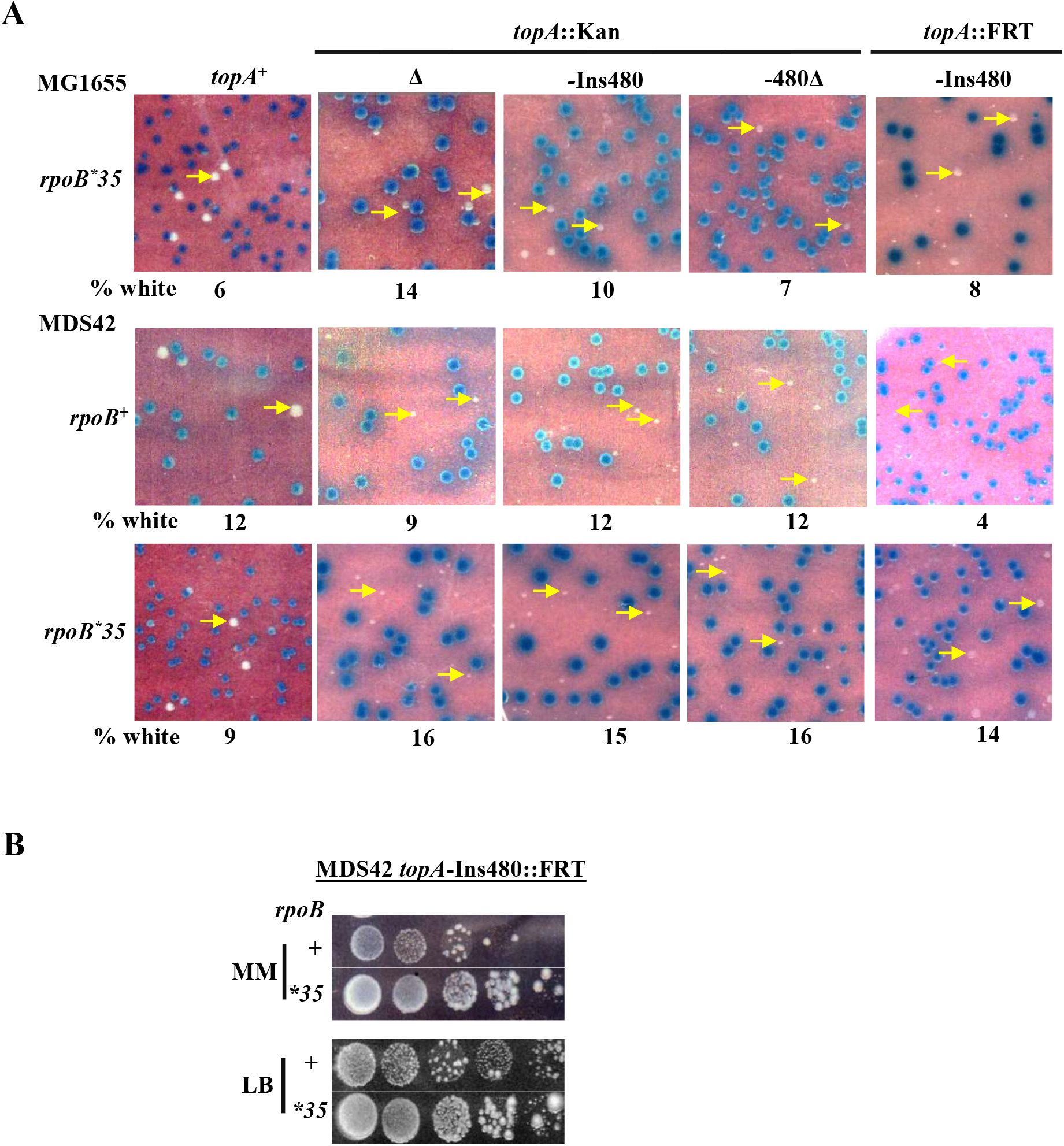
**(A)** Blue-white screening assay on glucose-minimal medium, of MG1655 or MDS42 strain derivatives with the *topA*^+^ shelter lasmid pHYD2390 and the different *topA* alleles as indicated on the top of each olumn; the *rpoB* allele status is indicated at left of each row. Examples of white olonies are marked by the yellow arrows. From left to right, strains used for the anels were pHYD2390 derivatives of: row 1, GJ16703, GJ16813, GJ16814, J16815 and GJ16854; row 2, GJ12134, GJ16816, GJ16817, GJ16818 and J 18977; and row 3, GJ16819, GJ16820, GJ16821, GJ16822 and GJ17777. (**B)** Serial dilution-spotting of *rpoB*^+^ and *rpoB*35* derivatives of MDS42 *topA-*ns480::FRT (GJ18977 and GJ17777, respectively) on LB and MM.

**Supplementary Figure S2.**
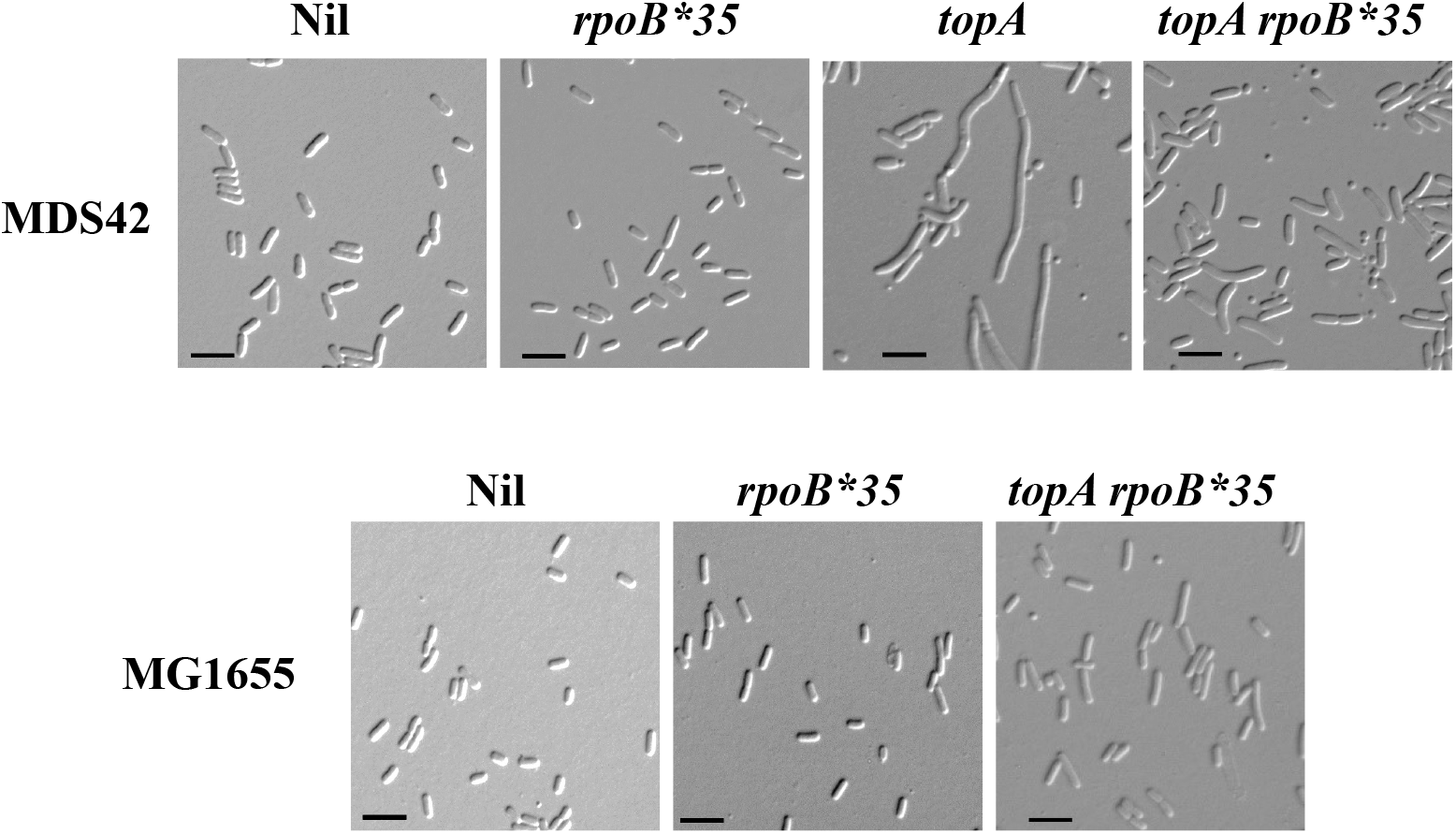
Cell morphology in derivatives of MDS42 and MG1655 (top and bottom rows, respectively), visualized by differential interference contrast microscopy. Scale bar (at bottom left of each panel), 5 µm. Relevant mutations in the strains are indicated on top of the corresponding panels. The *topA* allele in MDS42 derivatives was Δ*topA*::Kan and that in the MG1655 derivative was *topA-*Ins480::Kan. Strains used were (from left to right): top row, GJ12134, GJ16819, GJ16816, and GJ16820; and bottom row, GJ13519, GJ16703, and GJ16814.

**Supplementary Figure S3.**
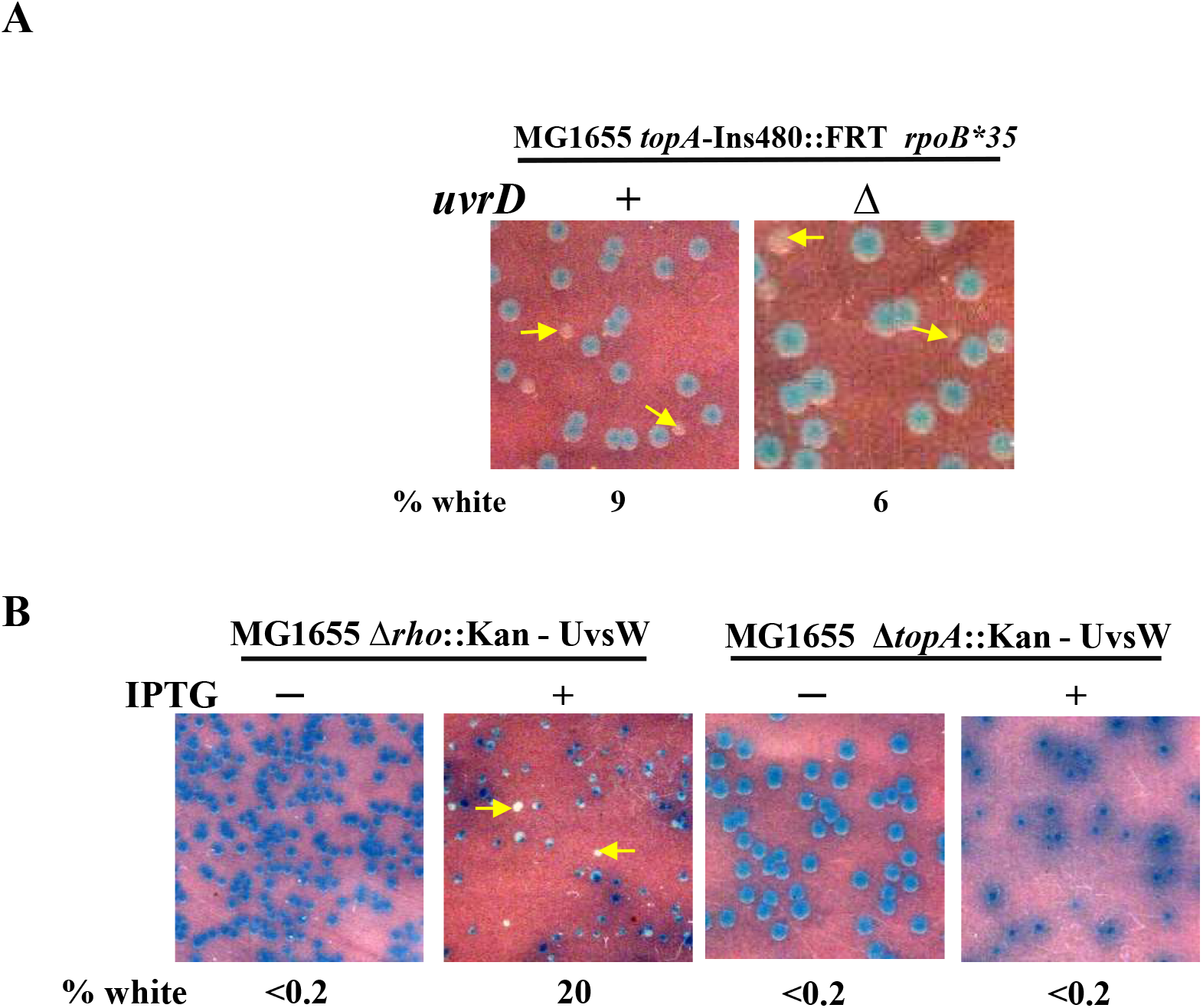
Blue-white screening assays with MG1655 derivatives (carrying cognate shelter plasmids) on glucose-minimal A medium to demonstrate that *topA* lethality rescue by *rpoB*35* is UvrD-independent **(A)**, and hat UvsW expression rescues Δ*rho* lethality but not Δ*topA* lethality **(B)**. Relevant genotypes are indicated on top of each of the panels. Examples of white colonies are marked by the yellow arrows. UvsW expression from a chromosomal P_tac_-UvsW construct was regulated by the omission (−) or addition (+) of IPTG (latter at 100 μM and 40 μM, respectively, for the *rho* and *topA* mutants). Strains used were: **(A)** pHYD2390 derivatives of GJ16854 (*uvrD*^+^) and GJ16880 (Δ*uvrD*); and **(B)** pHYD2411 derivative of GJ13531 (Δ*rho*) and pHYD2390 derivative of GJ15697 (Δ*topA*).

**Supplementary Figure S4.**
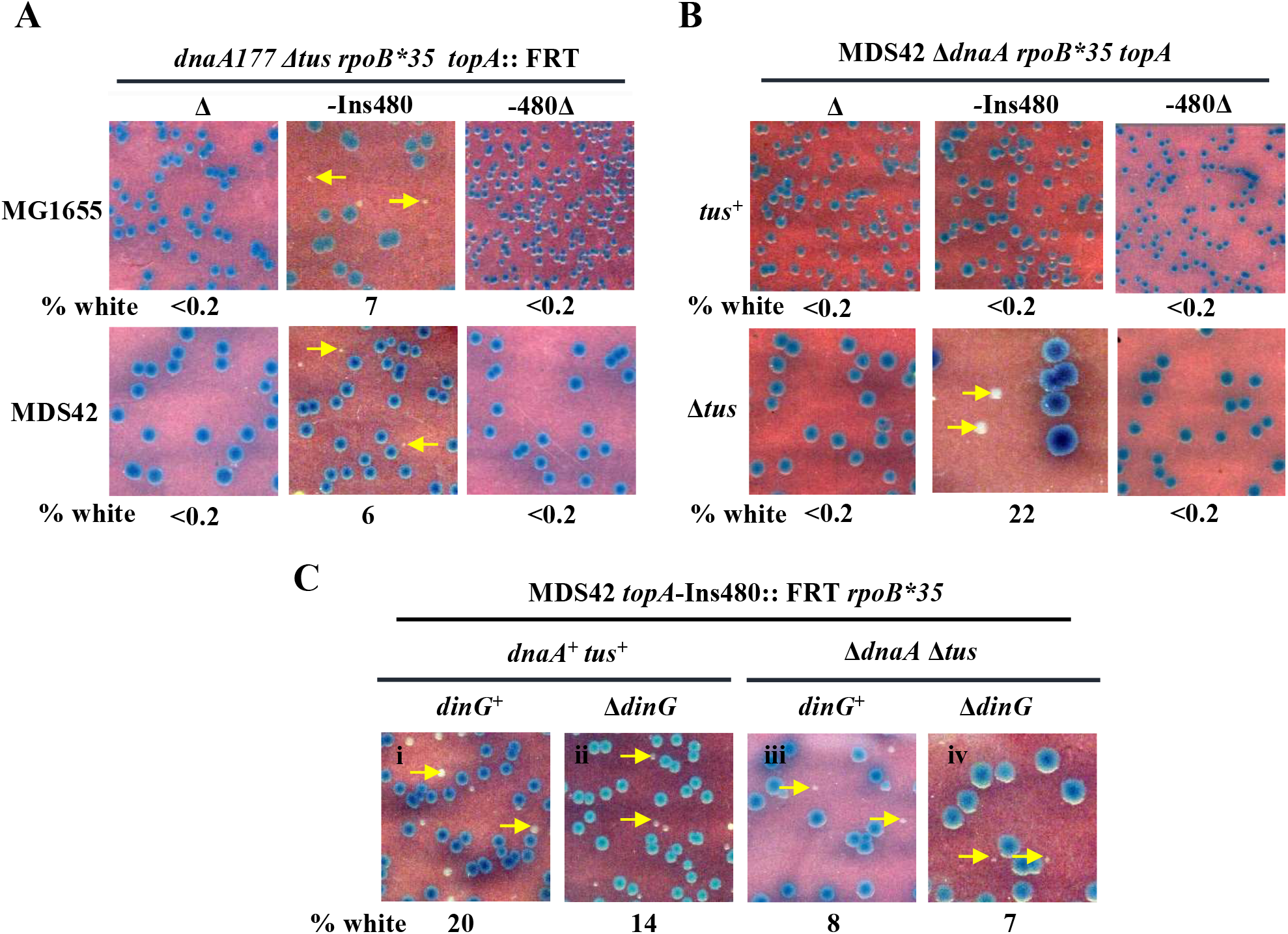
Features of *dnaA* lethality rescue by *topA* alleles. **(A-C)** Blue-white screening assays were performed at 30° on glucose-minimal A with *pA*^+^ *dnaA*^+^ shelter plasmid pHYD2390 in strain derivatives of MG1655 or DS42 whose relevant genotypes are indicated on top of each of the panels. xamples of white colonies are marked by the yellow arrows. From left to right, rains employed were pHYD2390 derivatives of: **A** top row, GJ18997, GJ15630 and GJ18999; **A** bottom row, GJ16902, GJ16903 and GJ16904; **B** top row, J18904, GJ18905 and GJ18906; **B** bottom row, GJ18907, GJ18908 and J18909; and **C**, GJ18918, GJ18921, GJ18941 and GJ18952.

**Supplementary Figure S5.**
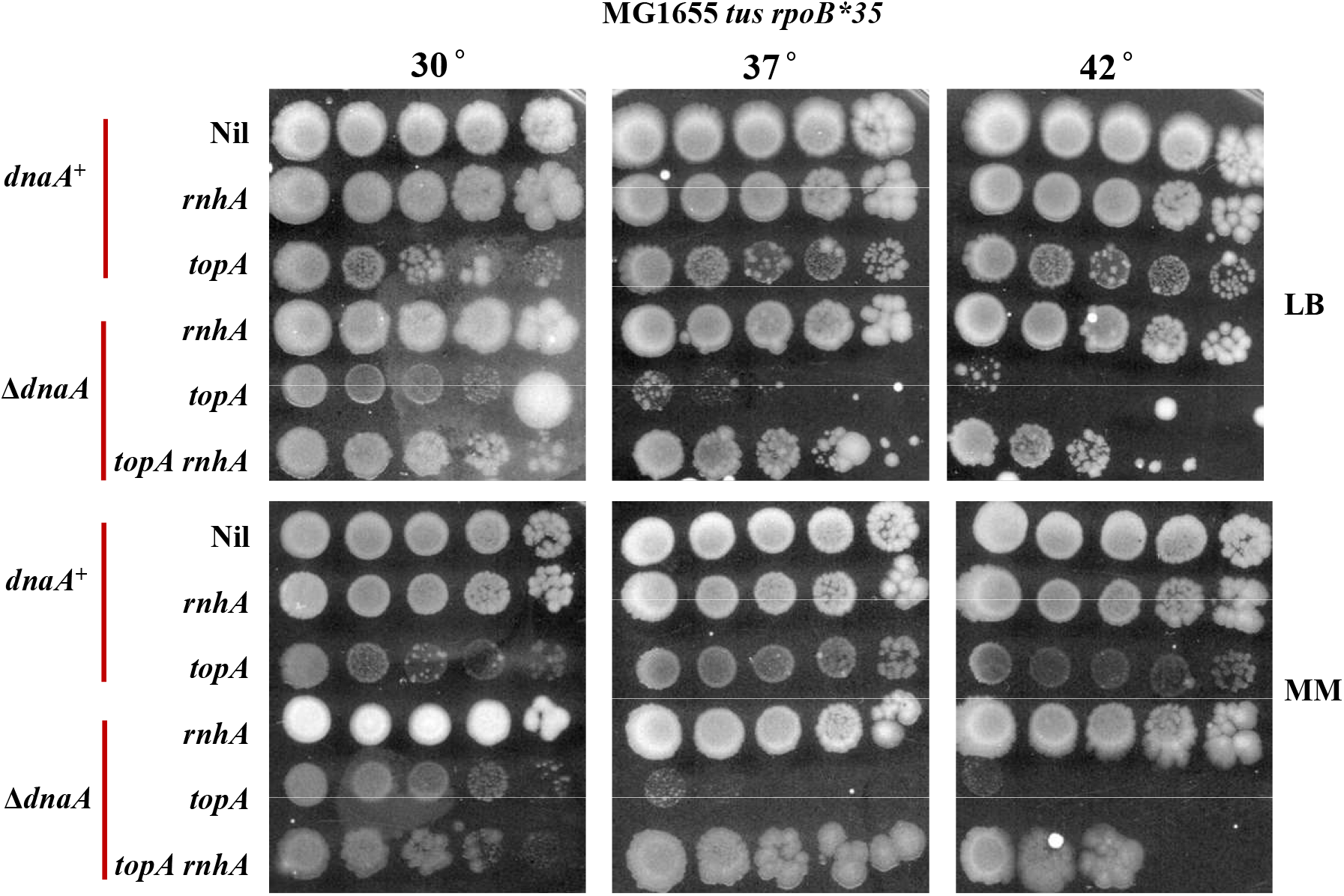
Serial dilution-spotting on LB and glucose-minimal A (MM) at indicated temperatures of derivatives of MG1655 Δ*tus rpoB*35* with additional relevant alleles as indicated at left. The Δ*dnaA*, *topA* and *rnhA* alleles were, respectively, Δ*dnaA*::FRT, *topA-*Ins480::FRT, and Δ*rnhA*::Kan. The six strains used were (from top): GJ17784/pHYD2390; GJ16475/pHYD2388; GJ17784; GJ16475; GJ18941; and GJ18951.

**Supplementary Figure S6.**
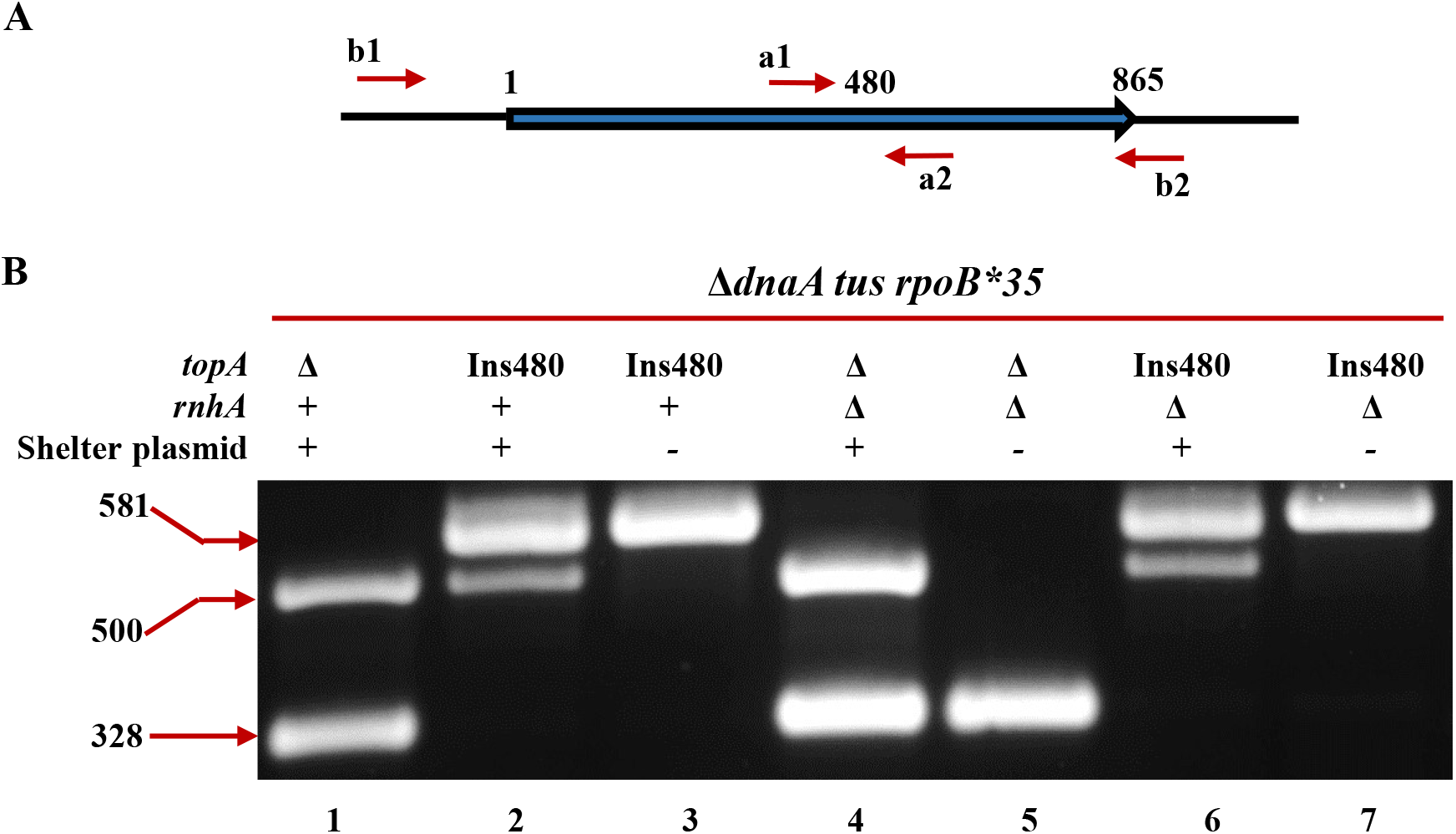
PCR validation of chromosomal *topA* genotype in Δ*dnaA* derivatives without or with *rnhA* mutation. **(A)** Schematic depiction of *opA* ORF (865 codons); positions of PCR primer pairs a1-a2 (5’-TGTACCAGTTAATCTGGCGTCAG-3’ and 5’-TCGTGATTTGCCACCTGGTCGAG-3’, respectively) flanking the codon 480 region, and b1-b2 (sequences given in Supplementary Methods above) for the entire ORF, are marked. The former primer pair is expected to yield amplicons of size 500 bp and 581 bp from *topA*^+^ and *topA-*Ins480::FRT, respectively, and no amplicon from Δ*topA*::FRT. The latter primer pair is expected to yield amplicons of size 2.8 kb and 328 bp from *topA*^+^ and Δ*topA*::FRT, respectively; however, the conditions used for PCR amplification were such as not to yield the large-size amplicons such as of 2.8 kb length. **(B)** Amplicons detected by gel electrophoresis after PCR with both primer pairs together, on DNA preparations from strains of genotype indicated on top of each ane. The *topA*^+^ *dnaA*^+^ shelter plasmid was pHYD2390. Allele designations used for *topA*: Δ, Δ*topA*::FRT and 480, *topA-*Ins480::FRT; and for *rnhA*: Δ, Δ*rnhA*::Kan. Migration positions of fragments of length 328, 500 and 581 bp are marked. Strains employed for the different lanes were: 1, GJ18940/pHYD2390; 2, GJ18941/pHYD2390; 3, GJ18941; 4, GJ18983/pHYD2390; 5, GJ18983; 6, GJ18951/pHYD2390; and 7, GJ18951.

## Notes

### Competing Interest Statement

The authors have declared no competing interest.

### Summary of Updates

Supplemenatl data file has been appended

